# Efficient cell-wide mapping of mitochondria in electron microscopic volumes using webKnossos

**DOI:** 10.1101/2024.10.20.619271

**Authors:** Yi Jiang, Haoyu Wang, Kevin Boergens, Norman Rzepka, Fangfang Wang, Yunfeng Hua

## Abstract

Recent technical advances in volume electron microscopy (vEM) and artificial intelligence-assisted image processing have facilitated high throughput quantifications of cellular structures, such as mitochondria that are ubiquitous and morphologically diversified. A still often overlooked computational challenge is to assign cell identity to numerous mitochondrial instances, for which both mitochondrial and cell membrane contouring used to be required. Here, we present a vEM reconstruction procedure (called mito-SegEM) that utilizes virtual path-based annotation to assign automatically segmented mitochondrial instances at the cellular scale, therefore bypassing the requirement of membrane contouring. The embedded toolset in webKnossos (an open-source online annotation platform) is optimized for fast annotation, visualization, and proofreading of cellular organelle networks. We demonstrate broad applications of mito-SegEM on volumetric datasets from various tissues, including the brain, intestine, and testis, to achieve an accurate and efficient reconstruction of mitochondria in a use-dependent fashion.

## INTRODUCTION

Mitochondria are vital organelles for eukaryotic cells, producing the energy-rich compound ATP that is essential for cellular metabolism. *In situ*, mitochondria not only vary in shape, size, and dynamics according to cell type and bioenergy status but also form highly organized networks with subcellular compartment precision to meet local metabolic demands.^1-3^ Therefore, spatial mapping of mitochondrial networks provides crucial information about key aspects of mitochondrial biology (such as morphological features, cellular positioning, and fission-fusion ratio) to comprehend the biological relevance of mitochondrial properties under healthy conditions or in disease states.^4^

Recent technical advances in volume electron microscopy (vEM) have enabled high-throughput 3D imaging of large tissue blocks at nanometer resolutions.^5-7^ Using this approach, the mitochondrial network organization has been profiled at an unprecedented scale in various cell types, including neurons,^8-10^ sensory cells,^11,12^ muscles,^13-15^ hepatocytes,^16-18^ sperm,^19,20^ and tumor cells.^21^ Currently, despite substantial progress in automated analysis powered by deep learning (DL) algorithms,^22-26^ manual annotation and proofreading of hundreds to thousands of instances are required to ensure trustful quantification of mitochondria, likely owing to the complexity and diversity of their morphology as well as prevalent inter-organelle contacts. Moreover, different sample preparations and imaging settings, cell type-specific mitochondrial heterogeneity, as well as unseen disease-related phenotypes, lead to poor DL model generalization.^22^ Thus, most DL pipelines still rely on repeated cycles of use-dependent fine-tuning. Lastly, it is often of interest to investigate the cellular landscapes of mitochondria, in particular those of polarized cells such as neurons,^9^ meaning that all mitochondria belonging to the same host cell are sought to be assigned based on the membrane boundaries of its interior. For intricate tissue systems like the brain, however, cellular volume reconstruction can be more challenging in terms of computational and human labor costs than the mitochondrial segmentation itself,^27^ making a high payoff for such analysis only on *ad hoc* densely segmented datasets.

Here, we report the development of a semi-automated analysis procedure (called mito-SegEM) for cell-wide mitochondrial reconstruction in large-scale volume EM data. It involves mitochondrial segmentation by a pre-trained artificial neural network and instance assignment using the path-based annotation tool of webKnossos as an alternative to the cell membrane contouring, thus achieving a highly efficient mapping of the cellular mitochondrial network in various specimens including brain tissues.

## RESULTS

Based on the online 3D data annotation tool webKnossos,^28^ we developed embedded functions that are aimed at facilitating the reconstruction of cellular mitochondrial networks in a semi-automated manner. The workflow (Figure 1A) included (1) automated mitochondrial segmentation, (2) uploading of mitochondrial masks to webKnossos for visualization (Video S1), (3) manual interconnecting of cellular mitochondrial instances by virtual paths (Video S2), and (4) proofreading and downloading of the mitochondrial merges associated with host cells or subcellular compartments (Video S3). Particularly, to enable fast interactive picking-up of mitochondrial instances by humans and avoid redundant annotation, all 3D masks of mitochondria are allowed to be hidden, and then individual of them can be highlighted with pseudo-color upon clicking on the vEM data (demonstrated in Video S2). In addition, on each selected mitochondrion, an annotation node was placed, and interconnected nodes within the cell assembled a virtual path that allows fast indexing of associated mitochondrial merges. Finally, a new keyboard shortcut was implemented in webKnossos for processing all segments that have been picked up by the path of annotation. This tool enables a fast selection of all mitochondria associated with the currently active path and adds them to a new segment group with the meshes automatically loaded for easy handling and 3D rendering (Video S4).

**Figure 1.**
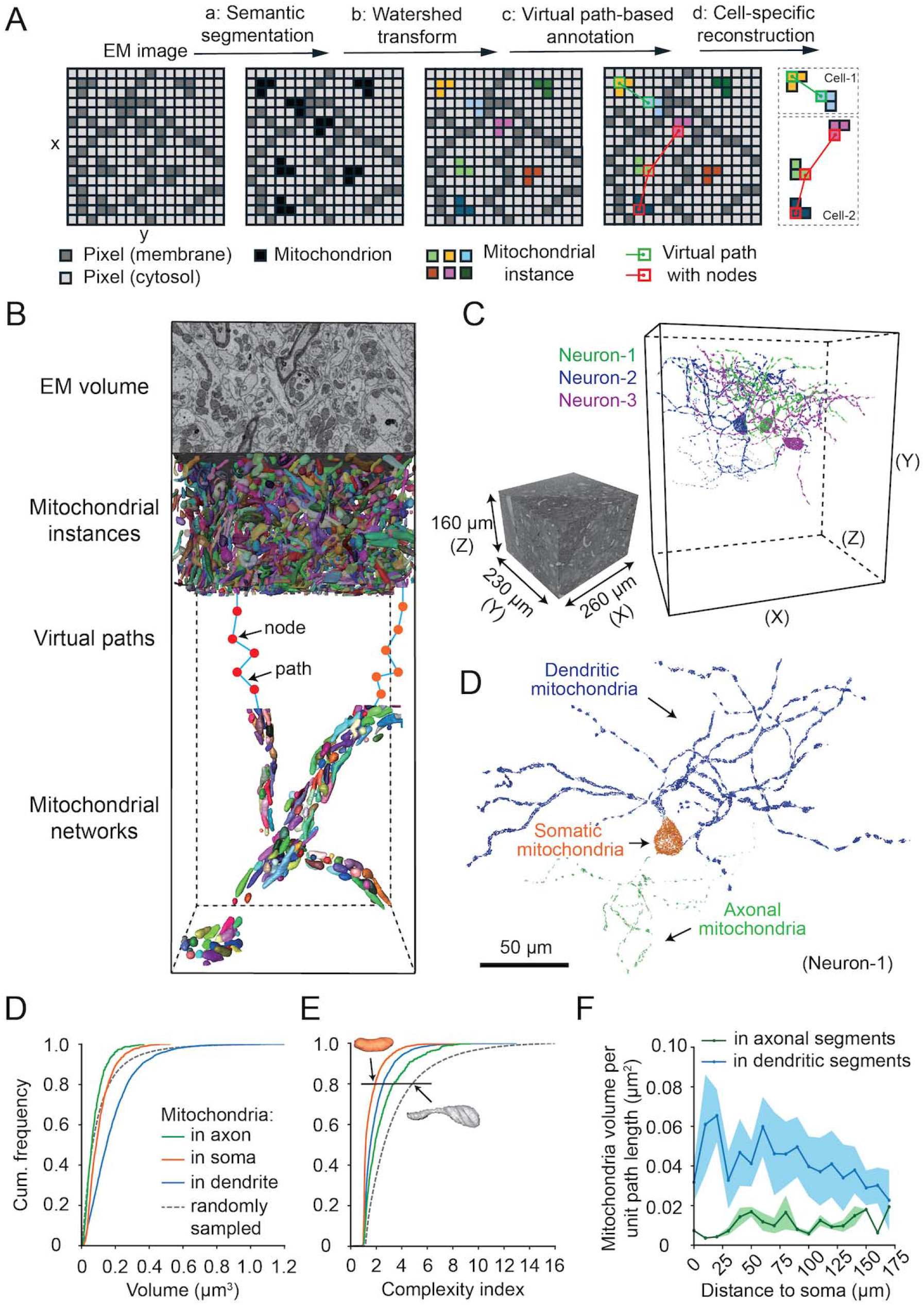
Reconstruction of the mitochondrial network in a neuron. (A) Flowchart of semi-automated analysis for cell-specific mitochondrial reconstruction using webKnossos. For the sake of simplicity, 2D pixel art is used as an example. (B) Schematic illustration of neuronal mitochondrial network reconstruction. (C) Dimensions of example SBEM volume acquired from the mouse brainstem (left). 3D rendering of the reconstructed mitochondrial networks (color-coded) from three neurons (right). (D) A representative cell-wide mitochondrial reconstruction of somatic (orange), dendritic (blue), and axonal (green) subpopulations of a neuron in C (Neuron-1). Scale bars, 50 µm. (E and F) Cumulative frequency distribution of mitochondrial instance volume and complexity index in the axons, dendrites, and soma of the neuron shown in D. (n = 650 axonal, n = 3,890 dendritic, n = 1,405 somatic mitochondria, n = 192,484 mitochondria in there randomly selected sub-volumes. See Figure S4 for details.) (G) In the same neuron as in D, dendrites (blue) and axons (green) were binned into 10 µm-long segments, in which mitochondrial volumes were measured, respectively. Data is presented as mean ± SD of mitochondrial volumes in segments at different path lengths away from the soma (See Online Methods for details).

### Mapping spatial organization of neuronal mitochondria

We first tested this procedure for neuronal mitochondrial reconstruction in the mouse brainstem tissue (Figure 1B). A dataset containing 3983 consecutive images with a pixel size of 15 × 15 nm^2^ was acquired using serial block-face scanning electron microscopy (SBEM)^29^ at a nominal thickness of 40 nm, yielding an EM volume of 260 × 230 × 155.7 µm^3^ after alignment along the cutting direction (Figure 1C). Automated segmentation of mitochondrial instances was performed using a pre-trained 3D U-net^25^ with dataset-specific minor modifications^30^ (Figure S1), which generated a map of 3D masks containing 1,048,575 mitochondria in the volume before proofreading. The EM volume together with the masks was then uploaded to webKnossos for the inspection and assignment of individual mitochondria to the corresponding host cells. For the proof-of-principle purpose, we randomly chose three neuronal cell bodies as starting points for neurite assignments of mitochondrial instances with our new tools.

The quantification revealed that each neuron accommodated 7495 ± 1146 (mean ± SD) mitochondrial instances, which was about 0.75% of all the mitochondria in the dataset (Figure 1C and S2). Next, in one neuron we subdivided the mitochondria into three subpopulations based on cellular positions, i.e. in the soma, dendrites, or axons (Figure 1D) by simply modifying the annotation paths in webKnossos. In contrast to random sampling, our cell-based quantification suggests neuronal compartment-specific differences in volume and complexity of mitochondria (Figure 1E and 1F). Note that the example cell features an overall higher ratio of simple mitochondria than those from randomly sampled volumes (Figure 1G), implying an inter-cellular heterogeneity in mitochondrial complexity. Furthermore, we showed distinct spatial distributions of mitochondria in the dendrites and axons (Figure 1H).

### Mitochondrial morphologies in mouse intestine and testis tissues

To highlight the general applicability of the mito-SegEM workflow, we analyzed two additional datasets from various mouse tissues. First, an intestinal sample block was cut at 50 nm step size, and 1000 consecutive slices were collected using automated tape-collecting ultramicrotome (ATUM).^31^ The SEM images were acquired at a pixel size of 8 × 8 nm^2^ and sequentially aligned to yield an EM volume of 41 × 57.3 × 50 µm^3^. Likewise, mitochondrial instances were automatically segmented and assigned to eight individual cells located at different regions of the intestinal crypt (Figure 2A and B). On average, each cell contained 80 ± 8 (mean ± SD) mitochondria (Figure S3A) with relatively broad size distributions (Figure 2C) and differential degrees of complexity (Figure 2D). Second, we reanalyzed a previously published SBEM dataset^32^ (dimension: 49.8 × 66 × 125 µm^3^, voxel size: 15 × 15 × 50 nm^3^) of mouse testis tissue (Figure 2E and F), in which twelve cells were randomly selected for mitochondrial reconstruction (491 ± 322, mean ± SD mitochondria per cell, Figure S3B). This revealed overall small-sized and simple mitochondria in the germ cells with a limited inter-cell difference (Figure 2G and H).

**Figure 2.**
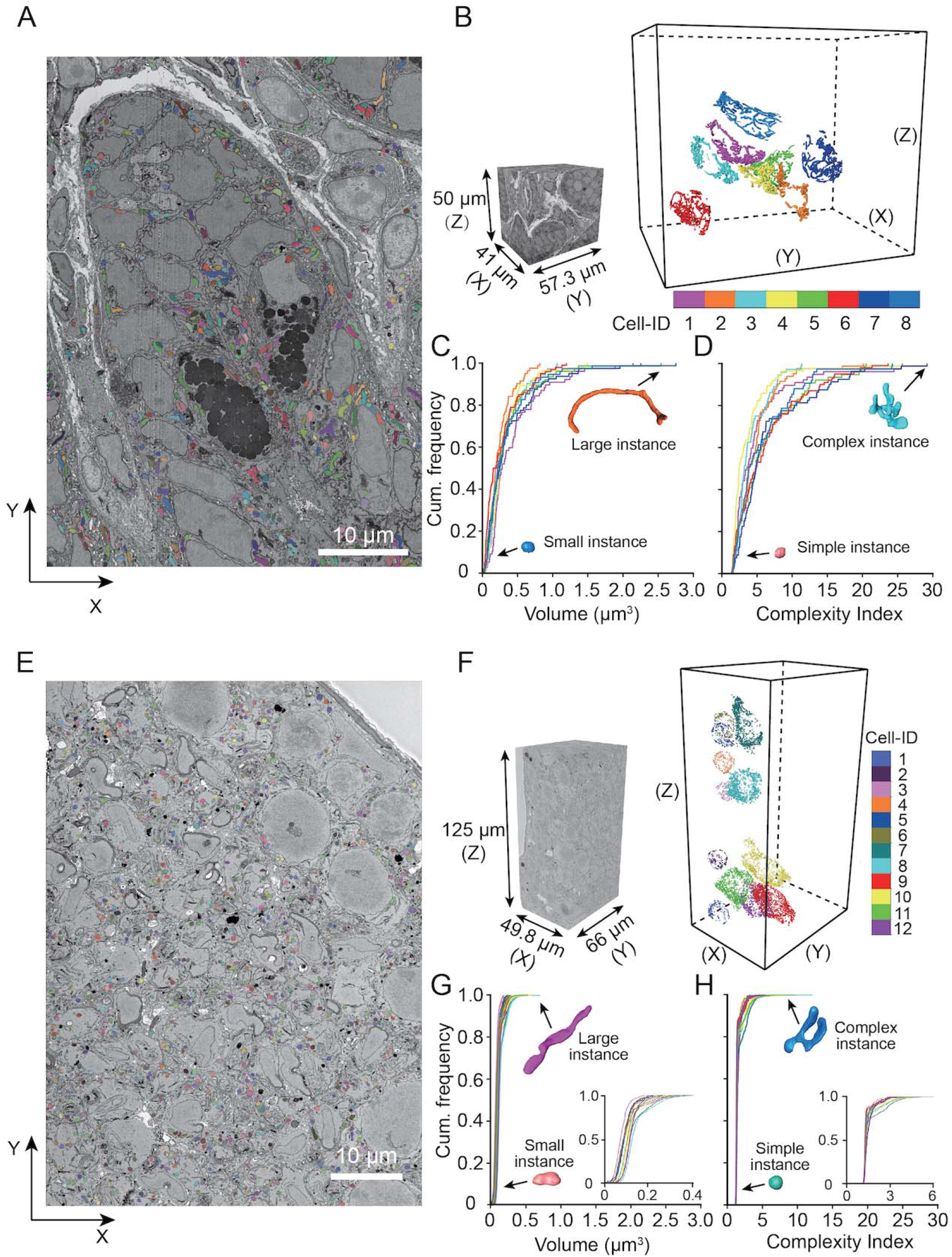
Cell-type specific differences in mitochondrial morphology. (A) EM image of a single ultra-thin slice from the mouse intestinal tissue. Mitochondria were automatically segmented and labeled with pseudo-colors. Scale bar 10 µm. (B) Volume of the acquired ATUM dataset (left), in which eight cells (color-coded) were randomly selected for mitochondrial reconstruction (right). (C and D) Cumulative frequency distributions of the volume and complexity index of mitochondrial instances per cell were computed from the reconstructions in B. (E) Example EM image with mitochondrial segmentations (pseudo-colored) of the mouse testis tissue block. Scale bar 10 µm. (F) SBEM volume (left) and 3D rendering of the mitochondrial reconstruction in twelve randomly selected cells. (G and H) Same as C and D, cumulative frequency distributions of mitochondrial instance volume and complexity index from the cellular reconstructions are shown in F. Insets: same plots but with an expanded x-axis.

### Annotation consumption for the mitochondrial assignment to host cells

Assigning individual mitochondria to corresponding host cells was an underestimated task that requires accurate segmentation of both mitochondrial and cell membranes to calculate their spatial overlapping. In the mito-SegEM workflow, we employed virtual path-based annotation to replace unnecessary contouring of the host cell membranes, whose annotation was usually done slice by slice. In neurons, for instance, contouring a small neurite fragment that contained one 1-μm-sized mitochondrion required manually tracing throughout 25 slices of 40-nm cutting thickness, while mito-SegEM simplified this to a single click on the mitochondrion. Thus, we speculated an increase in the annotation speed by one order of magnitude. To quantitatively compare the time costs between membrane contouring and virtual path-based annotation, we tested both approaches on the same datasets with pre-segmented mitochondria (Figure 3). For the dendrite fragment with a path length of 13.6 μm (Figure 3A-D), annotation of all 50 mitochondria using mito-SegEM was found 48.8 times faster than manual contouring of the membrane. Similarly, we reported a 21.8-fold reduction in the annotation consumption on a 13.3-μm-sized cell containing 77 mitochondria (Figure 3E-H). The varying degrees of annotation time savings are likely owing to different structural complexity and mitochondrial content between neurite and spherical cell.

**Figure 3.**
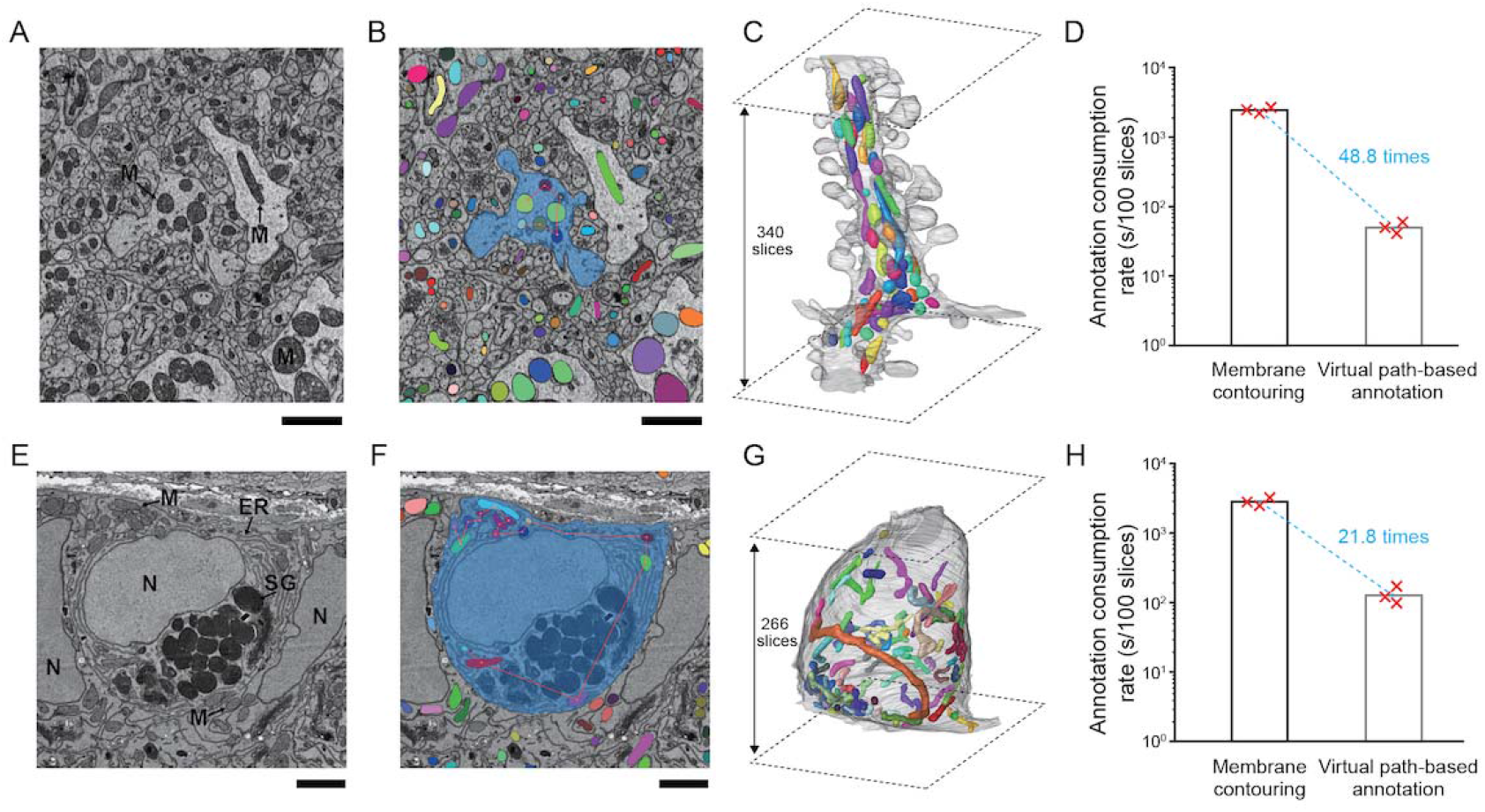
Annotation consumption needed for mitochondrial assignment. (A) Cropped EM image of the analyzed dendrite fragment from the mouse brainstem dataset. M: mitochondrion. Scale bar, 2 µm. (B) AI-segmented mitochondrial masks were labeled with pseudo-colors and overlaid on the raw image. Membrane contours of the dendrite (blue) were manually traced, within which annotated virtual paths were indicated as red lines. (C) The dendrite fragment (throughout 340 consecutive slices cut at a thickness of 40 nm) containing 50 mitochondria (colored) was rendered in 3D. (D) The normalized annotation durations required for slice-by-slice membrane contouring and annotation of mitochondria with virtual paths only. (E and F) An example cell was chosen from the mouse intestinal dataset. N: nucleus, ER: endoplasmic reticulum, M: mitochondrion, SG: secretory granule. The raw EM image (E) as well as its overlap (F) with mitochondrial masks (color-coded), the cell mask (blue), and virtual paths (red lines). Scale bar, 2 µm. (G) 3D rendering of the analyzed cell (throughout 266 consecutive slices, 50 nm thick) containing 77 mitochondria (colored). (H) Required annotation consumption for cell membrane contouring and virtual path-based mitochondrial annotation.

For cortical tissues, recently developed AI-based tools have enabled saturated segmentation, making the human annotation consumption rate about 25 times faster on neurite reconstruction.^33^ Although direct comparison was not performed, virtual path-based annotation seemed to be as fast as the focused annotation on AI-segmented neurites, without the need for computational resources and dataset-specific parameter tuning. Thereby, mito-SegEM is currently advantageous over other methods in terms of reconstruction costs in many scenarios and provides a fast alternative for studies whose primary goal is the mitochondrial network instead of dense neurite reconstruction.

## DISCUSSION

In the present work, we have developed mito-SegEM to facilitate use-dependent mitochondrial network reconstruction in electron microscopic volumes. By combining automatic mitochondrial segmentation and virtual path-based annotation, mito-SegEM enabled the efficient assignment of mitochondrial instances rather than individual pixels to the host cell. Besides, this procedure bypassed the contouring of the cell membrane borders that appear often intricated in tissues. We illustrated the broad applicability of our procedure on vEM datasets of the mouse brain containing intermingled neurites (Figure 1C), as well as the intestine and testis (Figure 2B and E), in which densely packed cells show less defined membrane borders.

Neurons are polarized cells with anatomically distinct compartments. This feature helps mitochondria maintain microenvironments in support of unique subcellular functions,^1,34^ including synaptic plasticity,^35-37^ axonal branching,^38,39^ and presynaptic release.^39-41^ Thus, it is of particular significance for both basic neuroscience and clinical studies to quantitatively characterize the compartment-specific mitochondrial morphologies in healthy individuals and disease models. While recent progress in vEM methods provides an unprecedented opportunity to visualize the cellular landscape of mitochondrial networks at nanometer resolution in 3D,^9^ a fast extraction of this information relies on existing full-volume reconstructions of neurites, to which individual mitochondria could be automatically assigned. Just like SegEM,^42^ our procedure, which profits from well-established neuronal mitochondrial segmentation and efficient skeleton annotation, can achieve presumably a 20-to-50-fold reconstruction efficiency gain (Figure 3D and 3H) and make a neurite-specific mapping of mitochondrial morphology affordable in most laboratories (Figure 1C). In addition, the skeleton-like annotation not only allows a real-time update of assigned mitochondria upon split and merge operations of neurites during proofreading in webKnossos, but also enables a spatial correlation of the mitochondrial network with numerous published neurite tracings and synapse annotations.^43-46^ Moreover, it has been recently shown that mitochondria can be used to facilitate neurite tracing in a focused annotation mode due to their elongation and ubiquitous presence in a neuron.^47^

Much progress has been made in improving the accuracy of mitochondrial segmentation using an adaptive template transformer (ATFormer) combined with a hierarchical attention learning mechanism on simultaneous identification of the mitochondrial mask and contour.^25,26^ Moreover, it has been recently demonstrated that promising inference results of mitochondrial segmentation can be readily delivered by a generalist DL model trained on massive EM datasets.^22^ However, given the structural diversity and different imaging settings, quantifications of mitochondrial morphologies without proofreading would not be granted in most case studies. In principle, the mito-SegEM procedure will allow transfer learning of a generalist model through human-in-the-loop annotation of mitochondria within the cells of interest only, bypassing the excessive requirement of model perfection for datasets with intermixed cell types or cells at different phases. Another advantage of this analysis routine is that the result from path annotation can be applied to the same dataset with mitochondrial masks inferred by different DL models, allowing the assignment of mitochondria with progressively improved segmentation outcomes. To enhance this function, further work will incorporate special nodes to interactively correct false split or merge errors.

Besides mitochondria, content-rich vEM datasets enable investigations of various organelles, for instance, endoplasmic reticula, lipid droplets, and lysosomes, as well as inter-organelle contact sites.^16,17,48,49^ We speculate that our analysis routine can be easily extended to a multi-organelle approach by parallelizing reconstructions of different organelles with specialist DL models, serving as a powerful tool for nanoscale mapping of a comprehensive cell atlas.

## Author Contributions

Y.H. conceived and supervised the study; Y.J. conducted the experiments with critical inputs from H.W. and K.M.B.; N.R. developed the software; F.W. acquired the EM datasets. Y.H. wrote the manuscript with contributions from all authors.

## Acknowledgments

We thank Drs. Hua Han and Linlin Li for the ATUM data acquisition. We thank the Transdisciplinary Platform of Functional Connectome and Brain-inspired Intelligence in Huairou Science City in Beijing and the Microscopic Technology & Analysis Center at the Institute of Automation, Chinese Academy of Sciences for providing technical support and device resources. This study is supported by the National Natural Science Foundation of China (82171133 to Y.H.), Industrial Support Fund of Huangpu District in Shanghai (XK2019011 to Y.H.), Innovative Research Team of High-level Local Universities in Shanghai (SHSMU-ZLCX20211700).

## STAR METHODS

## RESOURCE AVAILABILITY

### Lead Contact

Further information and requests for resources and reagents should be directed to and will be fulfilled by the Lead Contact, Y.H.

## Materials Availability

This study did not generate new unique reagents.

## Data and Code Availability

- Raw electron microscopy image data and segmentations supporting the current study are publically available via links listed in the key resources table.
- All original codes are publicly available on the publication date (https://github.com/JiangYi0311/Mito-SegEM).
- Any additional information required to reanalyze the data reported in this paper is available from the lead contact upon request.

## METHOD DETAILS

### Volume electron microscopy datasets

The mouse brainstem SBEM dataset contained 3983 single images (16000 × 16000 pixels) that were acquired at 15 nm pixel size and 40 nm cutting thickness using a field-emission scanning electron microscope (SEM, Gemini300, Carl Zeiss) equipped with an in-chamber ultramicrotome (3ViewXP, Gatan). The mouse intestine dataset was produced by SEM imaging on 50-nm-thick serial sections collected by a commercial ATUM setup (RMC Boeckeler, ATUMtome), yielding 1000 images of 6000 × 8000 pixels at 8 nm pixel size. The mouse testis dataset was previously reported,^32^ which was acquired using SBEM and contained 2500 50-nm-spaced single tiles of 10000 × 4000 15-nm-sized pixels.

### Image alignment and cubing

The alignment of both SBEM datasets (brainstem and testis) was performed offline using a script developed in our lab in MATLAB (MathWorks, US) to compute image offsets based on the cross-correlation maximum between consecutive sections.^45^ For the intestine dataset, a coarse-to-fine strategy was adopted to align the serial sections owing to nonlinear distortion. In detail, coarse alignment was performed by extracting the corresponding points between adjacent images using an affine transformation model, followed by fine alignment that involved pairwise correspondence extraction between adjacent images by SIFT flow,^50^ global adjustment of correspondence positions, and image wrapping through the moving-least-square method.^51^

After alignment, all datasets were cubed in 3D using a Python script included in the webKnossos toolkit (https://github.com/scalableminds/webknossos-libs) and uploaded to webKnossos for browsing and annotation.

### Automated mitochondrial segmentation

The pre-trained mitochondrial segmentation model^25,30^ was based on a residual 3D U-Net architecture with four-down/four-up layers (See Figure S1 for more details of the network architecture), which was provided by PyTorch Connectomics (https://connectomics.readthedocs.io/en/latest/). The model was trained to classify each voxel of the input stack (17 consecutive 256 × 256 pixel-sized images) into the “background”, “mitochondrial mask”, and “mitochondrial contour” categories. The model output was a two-channel image stack with the same format as the input, including the predicted probability maps of mitochondrial masks and contours. The overall loss function was

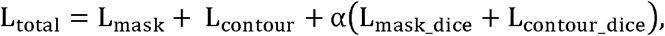

where L_mask_ and L_contour_ were the binary cross-entropy losses calculated for the mask and contour of mitochondrial segmentation, respectively. L_mask_dice_ and L_contour_dice_ were the Dice losses calculated for the mask and contour segmentation of mitochondria, respectively. α was a constant value and was set to 0.5.

### Model training and evaluation

The PyTorch deep learning framework (https://pytorch.org/) was employed for model training. The training datasets were randomly cropped from the raw image stacks. For the brainstem dataset three sub-volumes (1000 × 1000 × 100 voxels); for the intestine dataset four sub-volumes (one 1000 × 1000 × 100 voxels and three 2000 × 2000 × 100 voxels); and for the testis dataset two sub-volumes (1500 × 1500 × 100 voxels) were used. The ground-truth labels of mitochondria for the training datasets were generated by human experts using the Fiji^52^ plugin trakEM2^53^. For model training, the training datasets were pre-processed by padding and cutting pixel-by-pixel into small patches (256 × 256 × 17 voxels), which were randomly selected as inputs to conduct data augmentation, including rotation, flip, motion blur, noise addition, small region missing, and section missing. In addition, batch normalization was carried out during training, and the batch size was set to 4 due to hardware limitations. The model was trained for 300,000 iterations with a base learning rate of 0.04 using WarmupCosineLR (https://github.com/facebookresearch/detectron2) and asynchronous stochastic gradient descent on a single NVIDIA TITAN RTX GPU.

IOU and F1-score were used for model evaluation on three test datasets, which were cropped randomly from the raw data and labeled manually. The model-predicted values and human ground truth were compared, yielding an IOU of 0.88 and an F1-score of 0.94 for the test dataset of the brainstem. For the intestine and testis cases, an IOU of 0.92 and an F1-score of 0.96 as well as an IOU of 0.90 and an F1-score of 0.95 were obtained.

To generate mitochondrial instance masks, the seeds of mitochondria (or markers) were determined with a high mask probability and low contour probability by thresholding. Then, the marker-controlled watershed transform algorithm (part of the scikit-image library) was employed to generate high-quality instance masks of mitochondria with the seed locations and the predicted probability map of the masks.

### Mitochondria assignment by a virtual path

The segmentation of mitochondria was imported into the webKnossos using Python scripts. Next, in the “toggle merger mode” and with the option “hide the unmapped segmentation” selected, a start point was seeded and associated mitochondria were annotated one after another through the mouse right-clicks within individual instances. Upon each valid assignment, the corresponding mitochondrial instance would become visible with a pseudo-color and linked by an active node, so that missing and multiple annotations of mitochondrial instances could be minimized. Note that the “toggle merger mode” does not allow a mouse click outside the segments and ignores redundant annotations of a single segment. Finally, the assembly of the nodes was utilized to specify the associated mitochondrial instances that could be then operated as a defined group with i.e. self-written Python scripts.

### Quantitative analysis

Mitochondrial volume and surface area were calculated from the image stack using a Python script developed in our lab. The mitochondrial complexity index (MCI) was calculated using the formula^13^ as below:

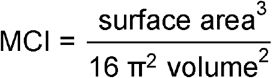

The skeleton for each neuron was split according to its compartment to generate individual sub-skeletons for the dendrites and axons. For each sub-skeleton, the nodes closest to the soma surface were chosen as the source nodes, and all other leaf nodes were marked as target nodes. The shortest path length from the source node to each target node was computed using Dijsktra’s algorithm. Each mitochondrion of the dendrite and axon was assigned to the closest skeleton node by calculating the distance between the mitochondrial centroid and the skeleton node. The mitochondrial volume per unit length was calculated by dividing the total mitochondrial volume within 10 µm of the dendritic or axonal path length by the 10 µm.

### Comparison of annotation consumption

In two separate annotation tasks, three independent human annotators were asked to trace the cell boundaries as well as annotate the mitochondria using the virtual path-based way for the same dendrite fragment (340 slices) and intestinal cell (266 slices) in webKnossos. The required durations could be directly read out from the saved annotation files containing the time points of each tracing or node placing (Tables S1 and S2).

## Referenc

1. Pekkurnaz, G., and Wang, X. (2022). Mitochondrial heterogeneity and homeostasis through the lens of a neuron. Nat Metab 4, 802–812. 10.1038/s42255-022-00594-w.

2. Giacomello, M., Pyakurel, A., Glytsou, C., and Scorrano, L. (2020). The cell biology of mitochondrial membrane dynamics. Nat Rev Mol Cell Biol 21, 204–224. 10.1038/s41580-020-0210-7.

3. Pernas, L., and Scorrano, L. (2016). Mito-Morphosis: Mitochondrial Fusion, Fission, and Cristae Remodeling as Key Mediators of Cellular Function. Annu Rev Physiol 78, 505–531. 10.1146/annurev-physiol-021115-105011.

4. Chan, D.C. (2020). Mitochondrial Dynamics and Its Involvement in Disease. Annu Rev Pathol 15, 235–259. 10.1146/annurev-pathmechdis-012419-032711.

5. Kievits, A.J., Lane, R., Carroll, E.C., and Hoogenboom, J.P. (2022). How innovations in methodology offer new prospects for volume electron microscopy. J Microsc-Oxford 287, 114–137. 10.1111/jmi.13134.

6. Titze, B., and Genoud, C. (2016). Volume scanning electron microscopy for imaging biological ultrastructure. Biol Cell 108, 307–323. 10.1111/boc.201600024.

7. Collinson, L.M., Bosch, C., Bullen, A., Burden, J.J., Carzaniga, R., Cheng, C., Darrow, M.C., Fletcher, G., Johnson, E., Narayan, K., et al. (2023). Volume EM: a quiet revolution takes shape. Nat Methods 20, 777–782. 10.1038/s41592-023-01861-8.

8. Faitg, J., Lacefield, C., Davey, T., White, K., Laws, R., Kosmidis, S., Reeve, A.K., Kandel, E.R., Vincent, A.E., and Picard, M. (2021). 3D neuronal mitochondrial morphology in axons, dendrites, and somata of the aging mouse hippocampus. Cell Rep 36, 109509. 10.1016/j.celrep.2021.109509.

9. Turner, N.L., Macrina, T., Bae, J.A., Yang, R., Wilson, A.M., Schneider-Mizell, C., Lee, K., Lu, R., Wu, J., Bodor, A.L., et al. (2022). Reconstruction of neocortex: Organelles, compartments, cells, circuits, and activity. Cell 185, 1082–1100 e1024. 10.1016/j.cell.2022.01.023.

10. Thomas, C.I., Keine, C., Okayama, S., Satterfield, R., Musgrove, M., Guerrero-Given, D., Kamasawa, N., and Young, S.M., Jr. (2019). Presynaptic Mitochondria Volume and Abundance Increase during Development of a High-Fidelity Synapse. J Neurosci 39, 7994–8012. 10.1523/JNEUROSCI.0363-19.2019.

11. Liu, J., Wang, S., Lu, Y., Wang, H., Wang, F., Qiu, M., Xie, Q., Han, H., and Hua, Y. (2022). Aligned Organization of Synapses and Mitochondria in Auditory Hair Cells. Neurosci Bull 38, 235–248. 10.1007/s12264-021-00801-w.

12. McQuate, A., Knecht, S., and Raible, D.W. (2023). Activity regulates a cell type-specific mitochondrial phenotype in zebrafish lateral line hair cells. Elife 12. 10.7554/eLife.80468.

13. Vincent, A.E., White, K., Davey, T., Philips, J., Ogden, R.T., Lawless, C., Warren, C., Hall, M.G., Ng, Y.S., Falkous, G., et al. (2019). Quantitative 3D Mapping of the Human Skeletal Muscle Mitochondrial Network. Cell Rep 26, 996–1009 e1004. 10.1016/j.celrep.2019.01.010.

14. Perez-Hernandez, M., Leo-Macias, A., Keegan, S., Jouni, M., Kim, J.C., Agullo-Pascual, E., Vermij, S., Zhang, M., Liang, F.X., Burridge, P., et al. (2021). Structural and Functional Characterization of a Na(v)1.5-Mitochondrial Couplon. Circ Res 128, 419–432. 10.1161/CIRCRESAHA.120.318239.

15. Bleck, C.K.E., Kim, Y., Willingham, T.B., and Glancy, B. (2018). Subcellular connectomic analyses of energy networks in striated muscle. Nat Commun 9, 5111. 10.1038/s41467-018-07676-y.

16. Parlakgul, G., Pang, S., Artico, L.L., Min, N., Cagampan, E., Villa, R., Goncalves, R.L.S., Lee, G.Y., Xu, C.S., Hotamisligil, G.S., and Arruda, A.P. (2024). Spatial mapping of hepatic ER and mitochondria architecture reveals zonated remodeling in fasting and obesity. Nat Commun 15, 3982. 10.1038/s41467-024-48272-7.

17. Jiang, Y., Li, L., Chen, X., Liu, J., Yuan, J., Xie, Q., and Han, H. (2021). Three-dimensional ATUM-SEM reconstruction and analysis of hepatic endoplasmic reticulum□organelle interactions. J Mol Cell Biol 13, 636–645. 10.1093/jmcb/mjab032.

18. Kang, S.W.S., Cunningham, R.P., Miller, C.B., Brown, L.A., Cultraro, C.M., Harned, A., Narayan, K., Hernandez, J., Jenkins, L.M., Lobanov, A., et al. (2024). A spatial map of hepatic mitochondria uncovers functional heterogeneity shaped by nutrient-sensing signaling. Nat Commun 15, 1799. 10.1038/s41467-024-45751-9.

19. Liu, P., Shi, J., Sheng, D., Lu, W., Guo, J., Gao, L., Wang, X., Wu, S., Feng, Y., Dong, D., et al. (2023). Mitopherogenesis, a form of mitochondria-specific ectocytosis, regulates sperm mitochondrial quantity and fertility. Nat Cell Biol 25, 1625–1636. 10.1038/s41556-023-01264-z.

20. Liu, J., Lin, L., Zhang, L., Ma, H., Chen, X., Pang, K., Li, L., and Han, H. (2024). Three-dimensional reconstruction of rat sperm using volume electron microscopy. Acta Biochim Biophys Sin (Shanghai) 56, 1699–1705. 10.3724/abbs.2024144.

21. Han, M., Bushong, E.A., Segawa, M., Tiard, A., Wong, A., Brady, M.R., Momcilovic, M., Wolf, D.M., Zhang, R., Petcherski, A., et al. (2023). Spatial mapping of mitochondrial networks and bioenergetics in lung cancer. Nature 615, 712–719. 10.1038/s41586-023-05793-3.

22. Conrad, R., and Narayan, K. (2023). Instance segmentation of mitochondria in electron microscopy images with a generalist deep learning model trained on a diverse dataset. Cell Syst 14, 58–71 e55. 10.1016/j.cels.2022.12.006.

23. Franco-Barranco, D., Lin, Z., Jang, W.D., Wang, X., Shen, Q., Yin, W., Fan, Y., Li, M., Chen, C., Xiong, Z., et al. (2023). Current Progress and Challenges in Large-Scale 3D Mitochondria Instance Segmentation. IEEE Trans Med Imaging 42, 3956–3971. 10.1109/TMI.2023.3320497.

24. Schubert, P.J., Dorkenwald, S., Januszewski, M., Jain, V., and Kornfeld, J. (2019). Learning cellular morphology with neural networks. Nat Commun 10, 2736. 10.1038/s41467-019-10836-3.

25. Wei, D., Lin, Z., Franco-Barranco, D., Wendt, N., Liu, X., Yin, W., Huang, X., Gupta, A., Jang, W.D., Wang, X., et al. (2020). MitoEM Dataset: Large-scale 3D Mitochondria Instance Segmentation from EM Images. Med Image Comput Comput Assist Interv 12265, 66–76. 10.1007/978-3-030-59722-1_7.

26. Pan, Y., Luo, N., Sun, R., Meng, M., Zhang, T., Xiong, Z., and Zhang, Y. (2023). Adaptive Template Transformer for Mitochondria Segmentation in Electron Microscopy Images. 2023 IEEE/CVF International Conference on Computer Vision (ICCV).

27. Motta, A., Schurr, M., Staffler, B., and Helmstaedter, M. (2019). Big data in nanoscale connectomics, and the greed for training labels. Curr Opin Neurobiol 55, 180–187. 10.1016/j.conb.2019.03.012.

28. Boergens, K.M., Berning, M., Bocklisch, T., Braunlein, D., Drawitsch, F., Frohnhofen, J., Herold, T., Otto, P., Rzepka, N., Werkmeister, T., et al. (2017). webKnossos: efficient online 3D data annotation for connectomics. Nat Methods 14, 691–694. 10.1038/nmeth.4331.

29. Denk, W., and Horstmann, H. (2004). Serial block-face scanning electron microscopy to reconstruct three-dimensional tissue nanostructure. PLoS Biol 2, e329. 10.1371/journal.pbio.0020329.

30. Lu, Y., Jiang, Y., Wang, F., Wu, H., and Hua, Y. (2024). Electron Microscopic Mapping of Mitochondrial Morphology in the Cochlear Nerve Fibers. J Assoc Res Otolaryngol 25, 341–354. 10.1007/s10162-024-00957-y.

31. Hayworth, K.J., Morgan, J.L., Schalek, R., Berger, D.R., Hildebrand, D.G., and Lichtman, J.W. (2014). Imaging ATUM ultrathin section libraries with WaferMapper: a multi-scale approach to EM reconstruction of neural circuits. Front Neural Circuits 8, 68. 10.3389/fncir.2014.00068.

32. Fan, Y., Huang, C., Chen, J., Chen, Y., Wang, Y., Yan, Z., Yu, W., Wu, H., Yang, Y., Nie, L., et al. (2022). Mutations in CCIN cause teratozoospermia and male infertility. Sci Bull (Beijing) 67, 2112–2123. 10.1016/j.scib.2022.09.026.

33. Motta, A., Berning, M., Boergens, K.M., Staffler, B., Beining, M., Loomba, S., Hennig, P., Wissler, H., and Helmstaedter, M. (2019). Dense connectomic reconstruction in layer 4 of the somatosensory cortex. Science 366. 10.1126/science.aay3134.

34. Alan, L., and Scorrano, L. (2022). Shaping fuel utilization by mitochondria. Curr Biol 32, R618–R623. 10.1016/j.cub.2022.05.006.

35. Kochan, S.M.V., Malo, M.C., Jevtic, M., Jahn-Kelleter, H.M., Wani, G.A., Ndoci, K., Perez-Revuelta, L., Gaedke, F., Schaffner, I., Lie, D.C., et al. (2024). Enhanced mitochondrial fusion during a critical period of synaptic plasticity in adult-born neurons. Neuron 112, 1997–2014 e1996. 10.1016/j.neuron.2024.03.013.

36. Rangaraju, V., Lauterbach, M., and Schuman, E.M. (2019). Spatially Stable Mitochondrial Compartments Fuel Local Translation during Plasticity. Cell 176, 73–84 e15. 10.1016/j.cell.2018.12.013.

37. Thomas, C.I., Ryan, M.A., Kamasawa, N., and Scholl, B. (2023). Postsynaptic mitochondria are positioned to support functional diversity of dendritic spines. Elife 12. 10.7554/eLife.89682.

38. Courchet, J., Lewis, T.L., Jr., Lee, S., Courchet, V., Liou, D.Y., Aizawa, S., and Polleux, F. (2013). Terminal axon branching is regulated by the LKB1-NUAK1 kinase pathway via presynaptic mitochondrial capture. Cell 153, 1510–1525. 10.1016/j.cell.2013.05.021.

39. Lewis, T.L., Jr., Kwon, S.K., Lee, A., Shaw, R., and Polleux, F. (2018). MFF-dependent mitochondrial fission regulates presynaptic release and axon branching by limiting axonal mitochondria size. Nat Commun 9, 5008. 10.1038/s41467-018-07416-2.

40. Ashrafi, G., de Juan-Sanz, J., Farrell, R.J., and Ryan, T.A. (2020). Molecular Tuning of the Axonal Mitochondrial Ca(2+) Uniporter Ensures Metabolic Flexibility of Neurotransmission. Neuron 105, 678–687 e675. 10.1016/j.neuron.2019.11.020.

41. Kwon, S.K., Sando, R., 3rd, Lewis, T.L., Hirabayashi, Y., Maximov, A., and Polleux, F. (2016). LKB1 Regulates Mitochondria-Dependent Presynaptic Calcium Clearance and Neurotransmitter Release Properties at Excitatory Synapses along Cortical Axons. PLoS Biol 14, e1002516. 10.1371/journal.pbio.1002516.

42. Berning, M., Boergens, K.M., and Helmstaedter, M. (2015). SegEM: Efficient Image Analysis for High-Resolution Connectomics. Neuron 87, 1193–1206. 10.1016/j.neuron.2015.09.003.

43. Schmidt, H., Gour, A., Straehle, J., Boergens, K.M., Brecht, M., and Helmstaedter, M. (2017). Axonal synapse sorting in medial entorhinal cortex. Nature 549, 469–475. 10.1038/nature24005.

44. Gour, A., Boergens, K.M., Heike, N., Hua, Y., Laserstein, P., Song, K., and Helmstaedter, M. (2021). Postnatal connectomic development of inhibition in mouse barrel cortex. Science 371. 10.1126/science.abb4534.

45. Hua, Y., Loomba, S., Pawlak, V., Voit, K.M., Laserstein, P., Boergens, K.M., Wallace, D.J., Kerr, J.N.D., and Helmstaedter, M. (2022). Connectomic analysis of thalamus-driven disinhibition in cortical layer 4. Cell Rep 41, 111476. 10.1016/j.celrep.2022.111476.

46. Karimi, A., Odenthal, J., Drawitsch, F., Boergens, K.M., and Helmstaedter, M. (2020). Cell-type specific innervation of cortical pyramidal cells at their apical dendrites. Elife 9. ARTN e46876 10.7554/eLife.46876.

47. Hong, B., Liu, J., Shen, L.J., Xie, Q.W., Yuan, J.B., Emrouznejad, A., and Han, H. (2023). Graph partitioning algorithms with biological connectivity decisions for neuron reconstruction in electron microscope volumes. Expert Syst Appl 222. ARTN 119776 10.1016/j.eswa.2023.119776.

48. Obara, C.J., Nixon-Abell, J., Moore, A.S., Riccio, F., Hoffman, D.P., Shtengel, G., Xu, C.S., Schaefer, K., Pasolli, H.A., Masson, J.B., et al. (2024). Motion of VAPB molecules reveals ER-mitochondria contact site subdomains. Nature 626, 169–176. 10.1038/s41586-023-06956-y.

49. Jang, W., Puchkov, D., Samso, P., Liang, Y., Nadler-Holly, M., Sigrist, S.J., Kintscher, U., Liu, F., Mamchaoui, K., Mouly, V., and Haucke, V. (2022). Endosomal lipid signaling reshapes the endoplasmic reticulum to control mitochondrial function. Science 378, eabq5209. 10.1126/science.abq5209.

50. Liu, C., Yuen, J., and Torralba, A. (2011). SIFT flow: dense correspondence across scenes and its applications. IEEE Trans Pattern Anal Mach Intell 33, 978–994. 10.1109/TPAMI.2010.147.

51. Schaefer, S., McPhail, T., and Warren, J. (2006). Image deformation using moving least squares. ACM Transactions on Graphics 25, 533–540. 10.1145/1141911.1141920.

52. Schindelin, J., Arganda-Carreras, I., Frise, E., Kaynig, V., Longair, M., Pietzsch, T., Preibisch, S., Rueden, C., Saalfeld, S., Schmid, B., et al. (2012). Fiji: an open-source platform for biological-image analysis. Nat Methods 9, 676–682. 10.1038/nmeth.2019.

53. Cardona, A., Saalfeld, S., Schindelin, J., Arganda-Carreras, I., Preibisch, S., Longair, M., Tomancak, P., Hartenstein, V., and Douglas, R.J. (2012). TrakEM2 software for neural circuit reconstruction. PLoS One 7, e38011. 10.1371/journal.pone.0038011.

